# Disease Tolerance Acquired Through Repeated *Plasmodium* Infection Involves Epigenetic Reprogramming of Innate Immune Cells

**DOI:** 10.1101/2023.04.19.537546

**Authors:** Jason Nideffer, Maureen Ty, Michele Donato, Rek John, Richard Kajubi, Xuhuai Ji, Holden Maecker, Felistas Nankya, Kenneth Musinguzi, Kathleen Dantzler Press, Bryan Greenhouse, Moses Kamya, Margaret E. Feeney, Grant Dorsey, PJ Utz, Bali Pulendran, Purvesh Khatri, Prasanna Jagannathan

## Abstract

The regulation of inflammation is a critical aspect of disease tolerance and naturally acquired immunity to malaria. Here, we demonstrate using RNA sequencing and epigenetic landscape profiling by cytometry by Time-Of-Flight (EpiTOF), that the regulation of inflammatory pathways during asymptomatic parasitemia occurs downstream of pathogen sensing—at the epigenetic level. The abundance of certain epigenetic markers (methylation of H3K27 and dimethylation of arginine residues) and decreased prevalence of histone variant H3.3 correlated with suppressed cytokine responses among monocytes of Ugandan children. Such an epigenetic signature was observed across diverse immune cell populations and not only characterized active asymptomatic parasitemia but also predicted long-term future disease tolerance when observed in uninfected children. This broad methylated signature likely develops gradually and was associated with age and recent parasite exposure. Our data support a model whereby exposure to *Plasmodium falciparum* induces epigenetic changes that regulate excessive inflammation and contribute to naturally acquired immunity to malaria.

## Introduction

The global disease burden of malaria is substantial. In 2020, there were an estimated 241 million cases of malaria, leading to 627,000 deaths worldwide^1^. The majority (77%) of these deaths were in children younger than the age of five^1^. The reason for this skewed distribution is not because older individuals have developed immunity that protects against *Plasmodium* infection^2^. Instead, these older individuals more frequently have low density infections that are asymptomatic, suggesting that repeated infection by *Plasmodium* conditions both an anti-parasite immune response and disease tolerance^3^.

Disease tolerance of malaria, the ability to sustain health despite considerable parasitemia^4,5^, is conceptually distinct but may partially rely on immunological tolerance, a state of non-reactivity towards antigens that normally would be expected to elicit an immune response^6^. For example, parasite-induced production of inflammatory cytokines by myeloid and other innate immune cells has been implicated in the pathogenesis of symptomatic (e.g. febrile) and severe malaria^7^. Therefore, attenuation of the innate pro-inflammatory response likely plays an important role in enabling asymptomatic infection. In support of this hypothesis, *ex vivo* transcriptional studies from malaria-naïve individuals undergoing experimental malaria infection have shown that malaria upregulates IFNγ, IL-1ß, and toll-like-receptor-mediated pro-inflammatory signaling and antigen presentation pathways^8,9^, with similar pathways upregulated in a study of Gabonese children with uncomplicated malaria^10^. These inflammatory responses were, however, blunted among a small subset of Malian adults, consistent with another study which found that prior malaria exposure may lead to dampening of genes involved in the type I and II interferon response^11^. Regarding the specific cellular populations which might be driving this modulated response, a recent report suggested that monocytes from malaria-exposed Malian adults produced lower levels of inflammatory cytokines IL-1ß, IL-6, and TNFα in response to *Plasmodium falciparum*-infected red blood cells (Pf-iRBC) compared with young Malian children^12^, suggesting that modified myeloid cell responses may facilitate disease tolerance.

Regarding mechanisms by which malaria may induce innate immune cell hyporesponsiveness to stimulation, it is increasingly recognized that innate cells can adapt following exposure to various antigenic stimuli—e.g., innate immune “memory.” For example, it has long been described that lipopolysaccharide (LPS) induces a durable state of monocyte cell refractoriness to subsequent LPS challenge^13^. More recently, stimulation with the tuberculosis vaccine bacilli Calmette-Guérin or β-glucans increases the long-term responsiveness of monocytes and/or bone marrow-derived myeloid precursors to microbial stimuli^14,15^. This modified responsiveness is thought to be mediated by changes in chromatin accessibility and cellular metabolism^16–18^, and has also been described for myeloid cells following vaccination against seasonal influenza^19^, as well as NK cells following CMV infection^20^. Two recent studies suggest that malaria antigen stimulation may modify the epigenetic state of monocytes, and that these changes may affect immune responses upon subsequent challenge^21,22^. However, *in vivo* evidence of malaria-induced epigenetic changes that modulate innate cell responsivity remains lacking.

In this study, we sought to determine whether immune cell epigenetics might contribute to disease tolerance of malaria in Ugandan children. We leveraged an unbiased, systems immunology approach using whole blood RNA sequencing, epigenetics by time of flight (EpiTOF)^23^, and an *in vitro* functional assay to demonstrate an association between patterns of histone methylation, asymptomatic parasitemia, and suppressed cytokine responses following pathogen recognition. We show that broad histone methylation among the immune cells of previously exposed Ugandan children predicts the acquisition of disease tolerance.

## Results

### Blood Transcriptomics Implicate Epigenetics in the Acquisition of Disease Tolerance

We performed RNA sequencing (RNA-seq) on paired, whole-blood samples from children living in malaria-endemic Uganda to determine how symptomatic malaria and asymptomatic parasitemia differentially affect the blood transcriptome compared with an uninfected baseline (Table 1). A differential expression analysis utilizing paired samples from the same infants (N=17 infants; N=34 samples) identified 2,779 genes that were upregulated and 2,307 genes that were downregulated during symptomatic malaria compared to an uninfected baseline. Notably, complement genes (*C1QA, C1QB, C1QC*), apoptotic genes (*PDL1*, *PDL2*, *APOL1*, *APOL2*, *APOL6*, *FAS*, *CASP7*), inflammatory caspases (*CASP1*, *CASP4*, *CASP5*), TLR pathway genes (*TLR1*, *TLR2*, *TLR4*, *TLR5*, *TLR6*, *TLR8*), cytokines (*IL6*, *IL10*), and granzyme (*GZMB*) were among the genes upregulated during symptomatic malaria (Figure 1A). Gene set enrichment analysis revealed interferon signaling and antigen processing/presentation as among the top 10 gene sets upregulated in children when they had symptomatic malaria (Figure 1B). In contrast, a separate experiment utilizing paired samples from children ages 2-12 (N=14) revealed no genes upregulated or downregulated compared to an uninfected baseline when children had asymptomatic parasitemia (Figure 1C), while gene sets related to phagocytosis, B cell and FC receptor signaling, complement activation, and calcium-mediated immune cell activation were among the top 10 downregulated gene sets in these children (Figure 1D). Interestingly, nearly all gene sets related to TLR signaling were upregulated during symptomatic and asymptomatic infection (Figure 1E), but cytokine signaling-related gene sets that were upregulated during symptomatic malaria were mostly downregulated during asymptomatic parasitemia (Figure 1F). These data suggest that disease tolerance is mediated by the regulation of inflammation downstream of pattern recognition, perhaps at the level of epigenetics. Leveraging our transcriptomic data, we observed several genes implicated in epigenetic control that were differentially expressed during symptomatic malaria, including those encoding methyltransferases that were downregulated. *EZH1*, a writer of histone H3 lysine 27 (H3K27) methylation, as well as *PRMT1, PRMT2*, and *PRMT7*, writers of arginine methylation, were among those genes (Figure 1G). Additionally, H3-3A and H3-3B, the genes encoding histone variant H3.3, were upregulated during symptomatic malaria (Figure 1G). This pattern of differential expression implicates the downregulation of H3K27 methylation and arginine dimethylation as well as the upregulation of H3.3 in the immune response that is associated with symptomatic malaria.

**Figure 1.**
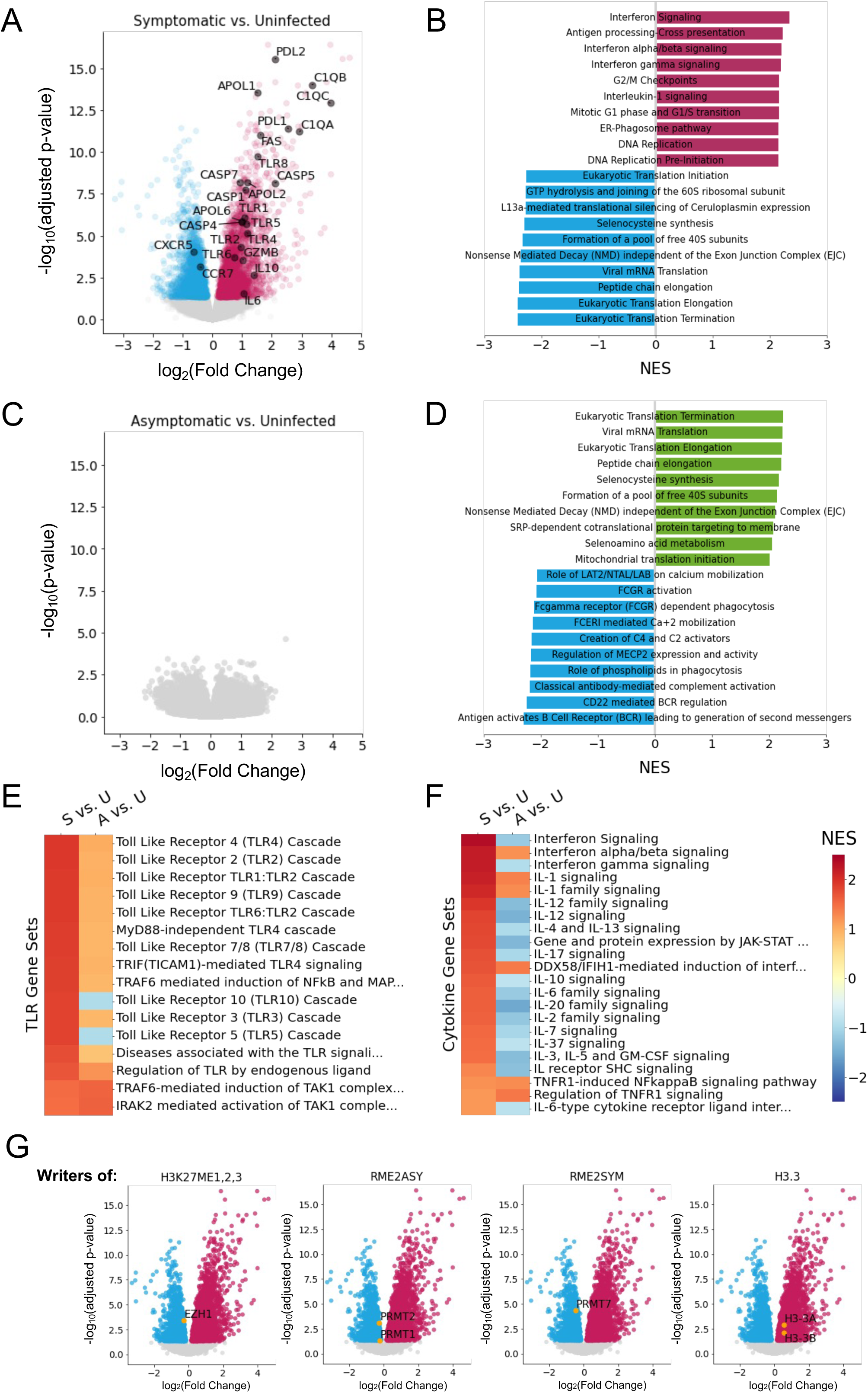
Differential gene expression in whole blood implicates epigenetics in the outcomes of disease during *Plasmodium* infection of Ugandan children. (A) Volcano plot showing differentially expressed genes in infants when they had symptomatic malaria (red) versus when they were uninfected (blue). P-values were adjusted to minimize false discovery. (B) Normalized enrichment scores (NES) for ten most enriched and ten most depleted Reactome gene sets in symptomatic versus uninfected. (C) Volcano plot showing differentially expressed genes in children when they had asymptomatic parasitemia (green) versus when they were uninfected (blue). P-values plotted were not adjusted because none were significant after adjustment. (D) NES for ten most enriched and ten most depleted Reactome gene sets in asymptomatic versus uninfected. (E-F) Enrichment of TLR signaling (E) or Cytokine related (F) gene sets in symptomatic malaria versus uninfected (S vs. U) and asymptomatic parasitemia versus uninfected (A vs. U). (G) Volcano plots highlighting differential gene expression of writers of histone markers in infants when they had symptomatic malaria versus when they were uninfected.

**Table 1.** Demographics and other characteristics of cohorts analyzed by whole-blood RNA sequencing.

| RNA Sequencing Cohorts |  |  |
| --- | --- | --- |
| **Symptomatic malaria cohort (n=17, paired)** |  |  |
|  | **Uninfected** | **Symptomatic** |
| Age in years, mean (SD) | 0.6 (0.11) | 0.6 (0.10) |
| Male gender, n (%) | 10 (58.8%) | 10 (58.8%) |
| Parasite density by blood smear, p/μl (range) | 0 | 39,000 (112-177000) |
| Sickle cell trait | 3 (17.7%) | 3 (17.7%) |
| **Asymptomatic parasitemia cohort (n=14, paired)** |  |  |
|  | **Uninfected** | **Asymptomatic** |
| Age in years, mean (SD) | 5.8 (2.8) | 5.9 (2.8) |
| Male gender, n (%) | 8 (57%) | 8 (57%) |
| Parasite density by qPCR, p/μl (range) | 0 | 918 (0.1-9035) |
| Sickle cell trait | 6 (42.8.0%) | 6 (42.8%) |

### Epigenetic Markers of Asymptomatic Parasitemia Correlate with Suppressed Cytokine Responses in Monocytes

To understand whether and how immune cell epigenetics contribute to disease tolerance, we performed epigenetic landscape profiling using EpiTOF on peripheral blood mononuclear cells (PBMCs) isolated from children living in malaria-endemic Tororo, Uganda (Figure 2A). The 12 children in this experiment (Table 2; separate from those studied by RNA-seq) were between the ages of 3 and 10 and had significant exposure to *P. falciparum* prior to their enrollment. Over the course of four years, these children were closely monitored for *Plasmodium* infections by repeated microscopy and loop-mediated isothermal amplication (LAMP) testing. Three PBMC samples from each child were analyzed by EpiTOF—one sample from when they were uninfected, one from when they had symptomatic malaria, and one from when they presented with asymptomatic parasitemia (Figure 2B). We assigned each of the 12 children to one of two cohorts (n=6 each), the samples from which were analyzed during two separate experiments (Figure 2A). By conducting the study in this way, we hoped to leverage inter-cohort heterogeneity resulting from reagent batch effects, procedural/instrument variability, and other sources of noise to arrive at generalizable and reproducible conclusions about the associations between immune cell epigenetics and disease tolerance.

**Figure 2.**
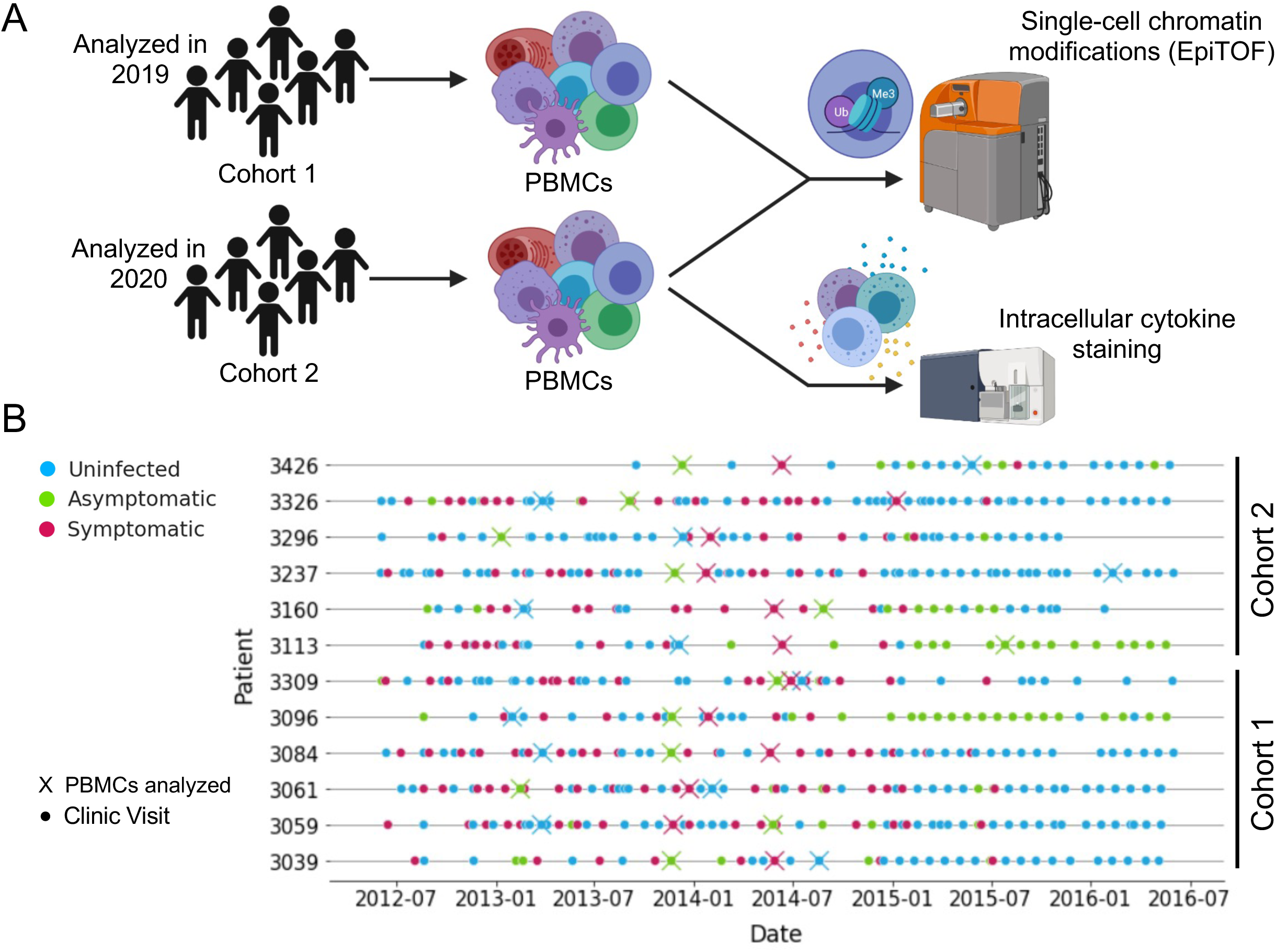
Experimental design to assess epigenetic signatures that characterize disease tolerance in children living in malaria-endemic Uganda. (A) Single-cell chromatin modifications were analyzed by EpiTOF. 12 children were sampled when they were uninfected, when they had asymptomatic parasitemia, and when they had symptomatic malaria. These children were split into two cohorts (n = 6), the samples of which were analyzed in two separate EpiTOF experiments performed in 2019 and 2020 for cohorts 1 and 2, respectively. Intracellular cytokine staining was also performed on the samples from cohort 2. (B) Timeline showing clinic visits (circles) and sample timepoints (X) of children included in our epigenetic study. Color indicates the disease state determined by qPCR and blood smear at the time of visit.

**Table 2.** Demographics and other characteristics of cohorts analyzed by EpiTOF.

| EpiTOF Cohorts |  |  |  |
| --- | --- | --- | --- |
| **Cohort 1 (n=6, paired)** |  |  |  |
|  | **Uninfected** | **Symptomatic** | **Asymptomatic** |
| Age in years, mean (SD) | 6.24 (0.78) | 6.64 (0.67) | 6.37 (1.03) |
| Male gender, n (%) | 3 (50%) | 3 (50%) | 3 (50%) |
| Parasite density by blood smear, p/μl (range) | 0 | 29000<br>(8160-56200) | 4539<br>(272-20000) |
| Sickle cell trait | 2 (33.3%) | 2 (33.3%) | 2 (33.3%) |
| **Cohort 2 (n=6, paired)** |  |  |  |
|  | **Uninfected** | **Symptomatic** | **Asymptomatic** |
| Age in years, mean (SD) | 6.67 (1.28) | 6.78 (1.50) | 6.50 (1.18) |
| Male gender, n (%) | 3 (50%) | 3 (50%) | 3 (50%) |
| Parasite density by blood smear, p/μl (range) | 0 | 9182<br>(336-30840) | 7787<br>(80-38880) |
| Sickle cell trait | 2 (33.3%) | 2 (33.3%) | 2 (33.3%) |

By examining the prevalence of epigenetic markers in surface marker-defined immune cell populations (Supplementary Figure 1), we identified classical and CD16+ monocytes as the most epigenetically distinct cell populations across disease states (Figure 3A). Among the markers differentially abundant in CD16+ monocytes was histone H3.3, which was less abundant in asymptomatic parasitemia versus symptomatic malaria (Figure 3A,B). As it is relevant, H3.3 has been implicated in the positive regulation of interferon stimulated genes in mice^24^. Methylation of H3K27 and H3K9, which has been associated with suppression of the interferon response^25^, was generally upregulated in CD16+ monocytes of children with asymptomatic parasitemia versus symptomatic malaria (Figure 3A,C). Finally, arginine dimethylation (*Rme2asy*, *Rme2sym*) was also more prevalent during asymptomatic infection (Figure 3A,D). This post-translational modification, in its asymmetric form, has primarily been shown to negatively regulate inflammation^26^. The differential regulation of these epigenetic markers across disease states is as would be predicted based on the differential expression of RNA encoding their writers (Figure 1G) and alludes to the possibility that the epigenetic state associated with asymptomatic infection may negatively regulate cytokine production.

**Figure 3.**
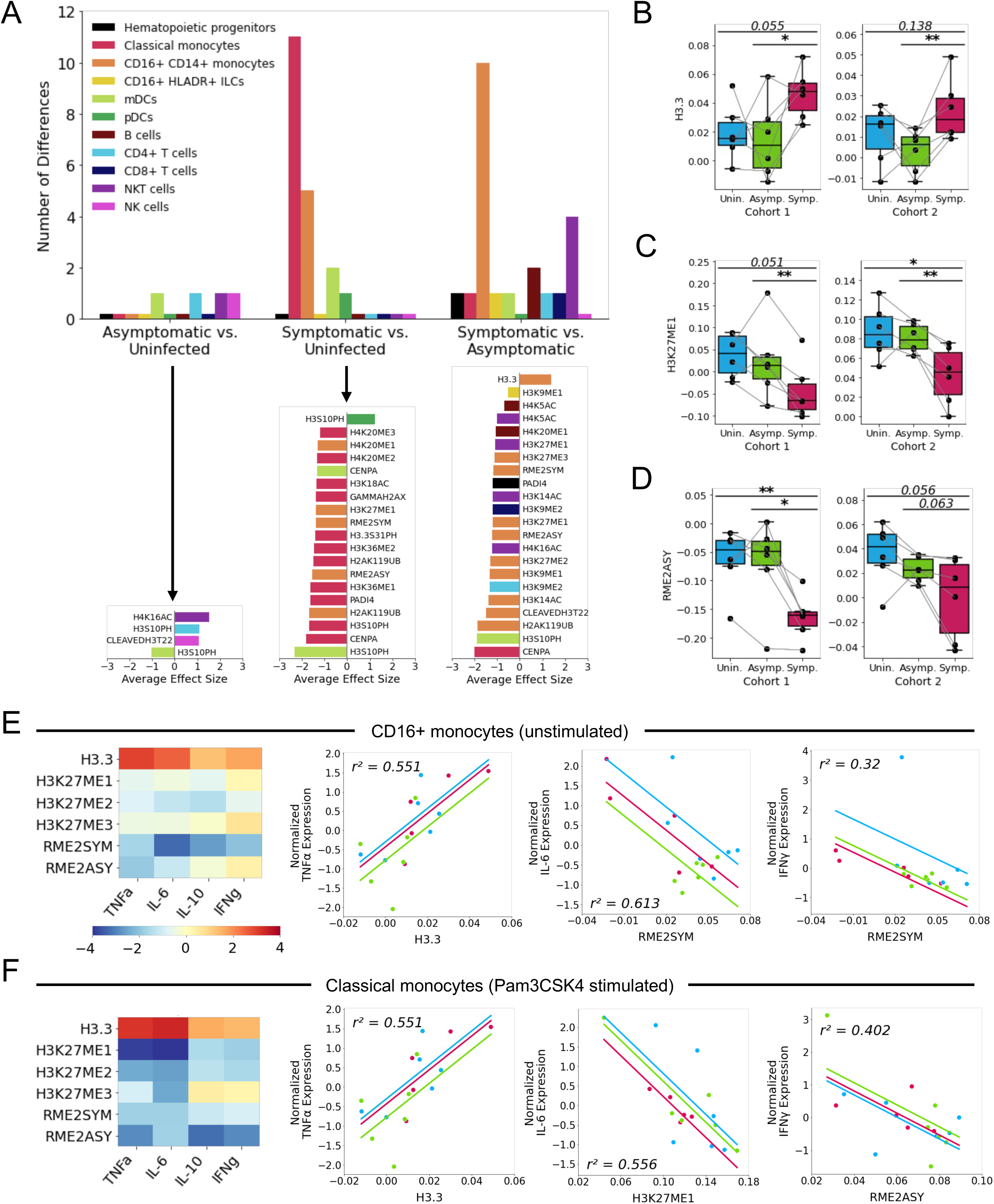
Epigenetic markers of asymptomatic parasitemia are associated with dampened cytokine responses in monocytes. (A) The number (top) and effect size (bottom; average of both cohorts) of differentially expressed epigenetic markers assessed via pairwise comparisons across disease states. Fisher’s combined probability test was used to combine p-values from both cohorts when effect size directions were consistent. Differentially expressed markers were defined as those that yielded a combined p-value of less than 0.01 to minimize false discovery. (B-D) Normalized expression of H3.3 (B), *H3K27me1* (C), and *Rme2asy* (D) in CD16+ CD14+ monocytes across disease states. (E-F) Heatmap displaying t-statistic values to describe how significantly the slope (cytokine expression regressed on histone marker abundance) deviates from zero (left side). Regressions were fit using random intercepts for each disease state to account for repeated measures. Select regressions are shown as scatterplots (right side; colored as in B-D according to disease state). Regressions demonstrate association between histone markers and cytokine expression in unstimulated CD16+ monocytes (E) and Pam3CSK4-stimulated classical monocytes (F). p-value < 0.05 (*); p-value < 0.01 (**).

We next performed flow cytometry with intracellular cytokine staining to examine PBMCs from the cohort 2 samples for the production of IFNγ, TNFα, IL-10, and IL-6 (Figure 2A, Supplementary Figure 2A). In doing so, we observed significant correlations between histone marker expression and basal cytokine production among CD16+ monocytes (Figure 3E). *H3.3* and *Rme2sym* were positively and negatively correlated (respectively) with the production of inflammatory cytokines such as TNFα, IL-6, and IFNγ (Figure 3E). CD16+ monocytes stimulated *in vitro* with the TLR2/TLR1 agonist Pam3CSK4 underwent considerable cell death (Supplementary Figure 2B), confounding our ability to study cytokine production by CD16+ monocytes following stimulation of this pathway. Pam3CSK4 stimulation did not, however, deplete classical monocytes and did strongly induce their expression of TNFα and IL-6 (Supplementary Figure 2C,D). Pam3CSK4-stimulated classical monocytes demonstrated strong correlations between epigenetics and cytokine responses consistent with those observed in CD16+ monocytes (Figure 3F). In sum, these data demonstrate that epigenetic patterns of asymptomatic parasitemia are associated with reduced inflammatory cytokine production by peripheral monocytes.

### An Immune Cell-Spanning Epigenetic Signature Predicts Disease Tolerance

While the associations between individual epigenetic markers and disease states/cellular functions are, themselves, novel, the epigenetic landscape of a cell is governed by the combined effects of diverse post-translational modifications^27^. Therefore, we performed unsupervised clustering of annotated PBMCs based on the abundance of epigenetic markers (excluding cell surface markers) in order to identify distinct epigenetic signatures spanning immune cell populations. This clustering, which was performed independently for cohorts 1 and 2, identified 17 epigenetic clusters in each cohort (Figure 4A). Meta clustering identified conserved epigenetic clusters (A-G) across both cohorts (Figure 4B). UMAP projections rotated and recolored according to meta clustering bore a striking resemblance (Figure 4C), suggesting reproducible clustering. In both cohorts, cluster G was enriched in children when they had asymptomatic parasitemia versus symptomatic malaria (Figure 4D,E). Interestingly, this cluster was defined by low H3.3 abundance but elevated H3K27 methylation (*H3K27me1*, *H3K27me2*, *H3K27me3*) and elevated arginine dimethylation (*Rme2sym*, *Rme2asy*) (Figure 4B), a pattern of expression that was observed in CD16+ monocytes during asymptomatic parasitemia and that was associated with diminished cytokine production (Figure 3). Cluster C, on the other hand, trended towards enrichment in symptomatic malaria compared to asymptomatic parasitemia (Figure 4D,F). This cluster was defined by very low methylation across several histone residues, including those that were abundantly methylated in cluster G (Figure 4B). Importantly, these epigenetic signatures were not only associated with symptomatic malaria (cluster C) and asymptomatic parasitemia (cluster G) in a heterogeneous population of PBMCs, but also demarcated more homogenous immune cell populations of monocytes and dendritic cells from children during symptomatic malaria versus asymptomatic parasitemia (Figure 4G). These results demonstrate that the broad adoption of an epigenetic signature—increased methylation (especially at H3K27 and arginine residues) and decreased H3.3—by diverse immune cells is associated with asymptomatic parasitemia.

**Figure 4.**
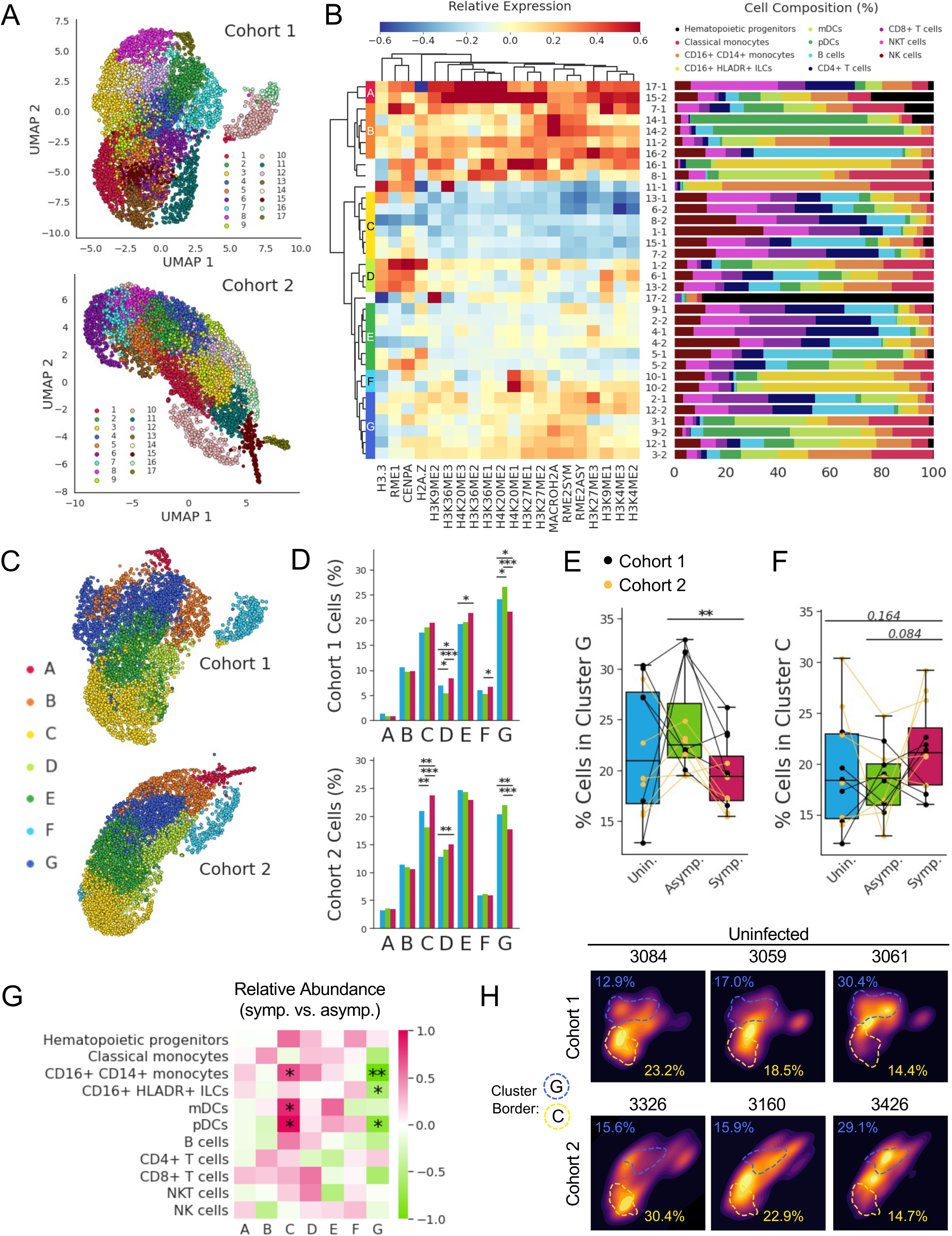
Symptomatic malaria and asymptomatic parasitemia are characterized by distinct and reproducible epigenetic signatures that span immune cell populations. (A) UMAP projections and Louvain clustering performed separately on single PBMCs from children of cohort 1 and cohort 2. (B) The heatmap (left) displays the relative expression of epigenetic markers (columns) across clusters (rows) from both cohorts. Original cluster identities are labeled with their cluster number and the cohort from which they were derived (i.e., cluster 17 from cohort 1 is denoted 17-1). Hierarchical clustering of the clusters yielded epigenetically-related meta clusters spanning both cohorts (labeled with letters A-G). The bar graph (right) displays the percent composition of each cluster by cell type. (C) Original UMAP projections of clusters 1 and 2 with cells recolored according to their meta cluster identity. The axis for the cohort 2 UMAP was rotated so that its orientation is consistent with the cohort 1 UMAP. (D) Percentage of cohort 1 (top) and cohort 2 (bottom) cells from a given disease state that were assigned to each meta cluster. (E-F) Percentage of each child’s clustered PBMCs assigned to cluster G (E) or cluster C (F). (G) Relative abundance of immune cell populations assigned to specific clusters when children from cohorts 1 and 2 (analyzed together) have symptomatic malaria versus when they have asymptomatic parasitemia. (H) Kernel density estimate plots depicting the distributions of cells across the UMAPs in ‘C’ for six different children at the uninfected timepoint. Dashed blue and yellow lines show the approximate borders of clusters G and C, respectively. Similarly colored percentages denote the proportion of each uninfected child’s cells that belong to the corresponding cluster. p-value < 0.05 (*); p-value < 0.01 (**); p-value < 0.001 (***).

We noticed that the distributions of cluster G and cluster C frequencies displayed greater variances among the uninfected timepoints compared to the timepoints of symptomatic malaria and asymptomatic parasitemia (Figure 4E). The epigenetic heterogeneity observed across uninfected children (Figure 4H), thus, led us to wonder whether a child’s baseline epigenetic state (when they are uninfected) primes them for a certain disease outcome upon infection. To answer this, we assessed the relationship between the frequencies of epigenetic clusters measured during an uninfected state with each child’s subsequent future risk of symptomatic malaria. Indeed, the percentages of cells within cluster G and cluster C among uninfected children were negatively and positively correlated (respectively) with their future incidence of symptomatic malaria (Figure 5A). However, the frequencies of these epigenetic clusters did not appear to affect future risk of parasitemia (Figure 5B) or future parasite burden (Figure 5C). Interestingly, cluster G and cluster C frequencies did predict future incidences of non-malarial fevers (Figure 5D), suggesting non-specific effects of epigenetic adaptations in these children that may have consequences beyond malaria.

**Figure 5.**
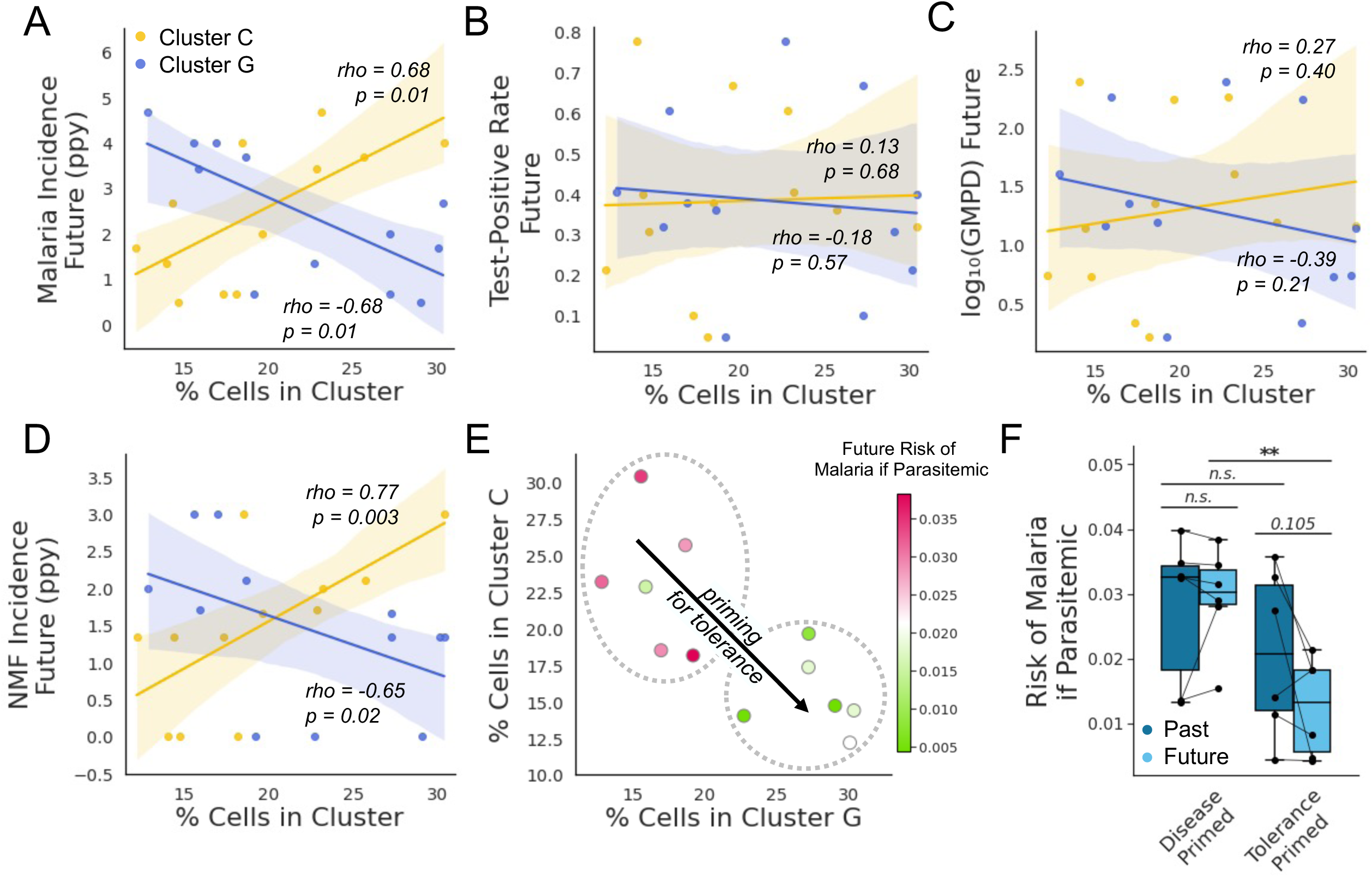
Immune cell epigenetics during homeostasis predict the acquisition of disease tolerance. (A-D) Spearman correlations between the percentage of an uninfected child’s sampled PBMCs assigned to cluster G (blue) and cluster C (yellow) versus their future incidence of malaria (A), their future test-positive rate (by blood smear) (B), their future parasite burden (log-scaled geometric mean parasite density [GMPD]) (C), and their future incidence of non-malarial fevers (NMF) (D). Future outcome variables were calculated using data collected for a duration of three years after the sample timepoint. Shaded regions depict 95% confidence intervals associated with a linear regression. (E) Scatterplot depicting the negative correlation between cluster G and cluster C frequencies. Each observation represents a different child, and its color denotes that child’s future risk of malaria if they test parasitemia-positive. Dashed circles highlight children that are “disease primed” (upper left) or “tolerance primed” (lower right). (F) Box plot depicting past and future risks of malaria given parasitemia. Children are stratified based on whether they are “disease primed” or tolerance primed (according to ‘E’). p-value < 0.01 (**).

Among uninfected children, cluster G and cluster C frequencies were negatively correlated (Figure 5E). Children with high cluster G but low cluster C frequencies had a much lower future risk of malaria given parasitemia and were considered “tolerance primed” (Figure 5E). Children with low cluster G but high cluster C frequencies were considered “disease primed.” Tolerance primed children, unlike disease primed children, had largely obtained disease tolerance and naturally acquired immunity, and they demonstrated a transition from a high-risk past to a low-risk future (Figure 5F).

### Trajectories of Epigenetic Reprogramming Model the Gradual Acquisition of Disease Tolerance Among Ugandan Children

Given the inverse relationship between cluster C and cluster G among children growing up in malaria-endemic Uganda, we wondered whether a trajectory might exist whereby over the course of childhood and through repeated *Plasmodium* exposure, immune cells gradually adopt a broad methylation signature (transitioning from cluster C to cluster G). To explore this idea, we performed trajectory inference using tSpace, which seeks to uncover pathways of gradual biological change among cells projected in “trajectory space,” where cells are defined in relation to one another by their position along nearest-neighbor pathways^28^. When applied to our EpiTOF data from uninfected children, this algorithm consistently yielded two trajectories (Figure 6A). One trajectory consisted of myeloid cells and the other consisted of lymphoid cells, but both trajectories demonstrated parallel epigenetic adaptations with the passage of pseudotime (Figure 6B). In this model, increasing pseudotime brought with it epigenetic changes associated with asymptomatic parasitemia and disease tolerance—decreasing H3.3 levels and increasing H3K27 methylation/arginine dimethylation (Figure 6C). Additionally, passage through pseudotime demonstrated clear changes in the abundance of cluster C (which was prevalent only early in pseudotime) and cluster G (which appeared later in pseudotime) (Figure 6C).

**Figure 6.**
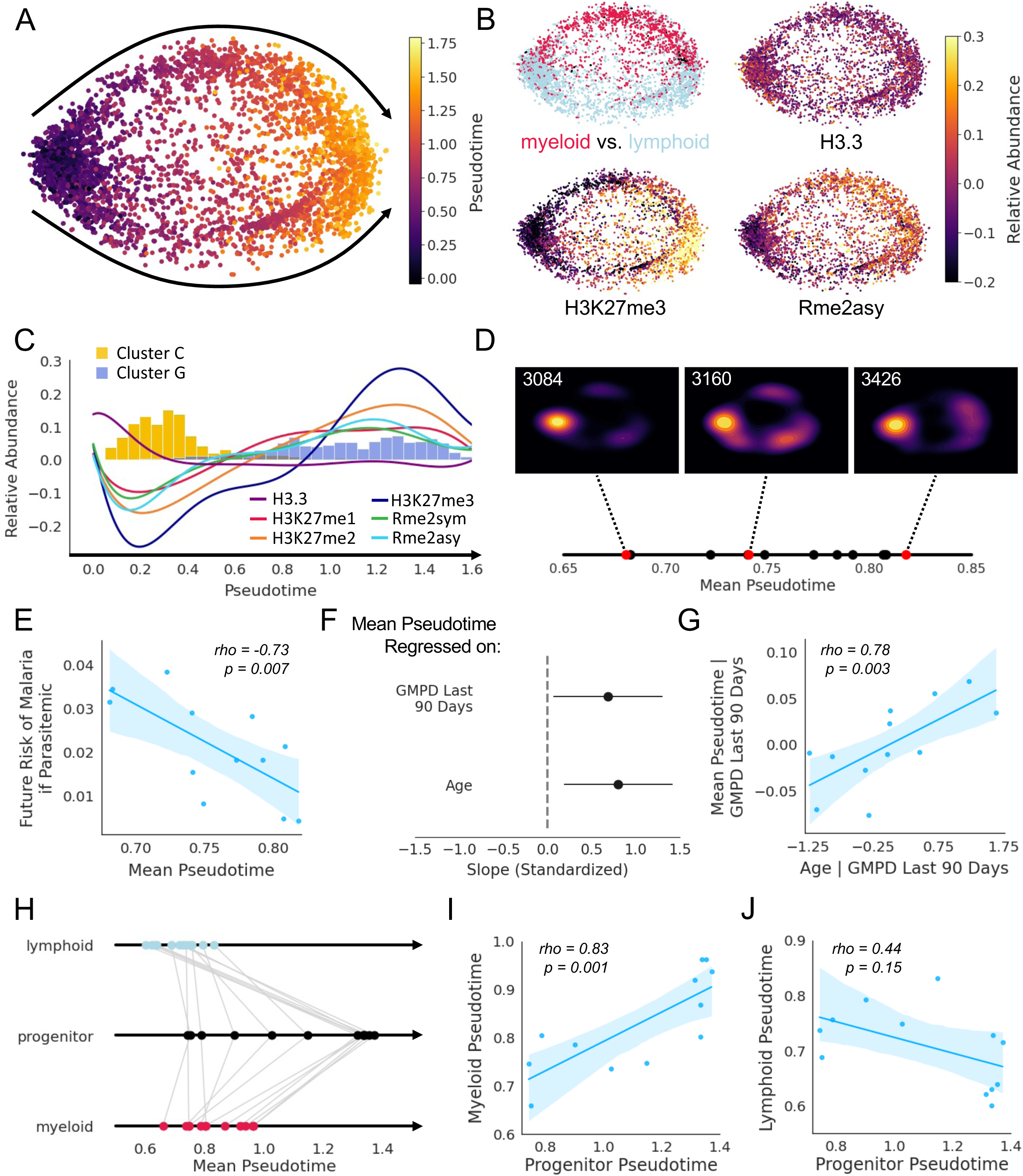
Pseudotime analyses model epigenetic changes that occur during childhood and contribute to disease tolerance among malaria-exposed Ugandans. (A) tSpace projections of cells from uninfected children analyzed by EpiTOF. Cells from both cohorts (depicted in Figure 4) were used and are colored based on pseudotime values. Black arrows depict two parallel trajectories through pseudotime. (B) tSpace projections colored by cell lineage (top left), H3.3 abundance (top right), *H3K27me3* abundance (bottom left), or *Rme2asy* (bottom right). (C) Epigenetic marker (colored lines) and cluster (colored histograms) abundance over pseudotime. (D) Mean pseudotime values calculated on a per-child basis are plotted along a solid line. Kernel density estimate plots are depicted for three children with early (3084), middle (3160), and late (3426) mean pseudotime values (red dots). (E) Correlation between mean pseudotime and future risk of malaria if parasitemic, demonstrating that children who have advanced further through pseudotime are more disease tolerant. (F) Fitted parameter values from a multiple regression of mean pseudotime on age and recent parasite burden (measured as geometric mean parasite density over the preceding 90 days)—values of fitted parameters are shown with 95% confidence intervals. Independent and dependent variables were standardized prior to model fitting. (G) Partial regression where the effect of recent parasite burden is removed and mean pseudotime is regressed on age. (H) Mean pseudotime calculated for lymphoid, progenitor, and myeloid cells. Gray lines connect values calculated from cells of the same child. (I) Correlation between myeloid pseudotime and progenitor pseudotime. (J) Correlation between lymphoid pseudotime and progenitor pseudotime. Shaded regions depict 95% confidence intervals associated with a linear regression.

Next, we investigated whether pseudotime in our model reflected chronological time in the lives of Ugandan children as they developed disease tolerance of malaria. To do this, we first calculated the mean pseudotime value for each of the twelve children (Figure 6D). A strong negative correlation between the mean pseudotime value and a child’s future risk of malaria if parasitemic was observed (Figure 6E), indicating that progression through pseudotime models not only a trajectory of epigenetic reprogramming but also a trajectory through which disease tolerance is achieved. Furthermore, a child’s age and recent parasite burden contributed to their progression along this trajectory (Figure 6F,G), suggesting that the epigenetic changes observed over pseudotime model the epigenetic changes that occur as Ugandan children grow up and are repeatedly infected with *Plasmodium*. Finally, we sought to determine whether epigenetic reprogramming within the myeloid compartment might be driven by epigenetic modifications to hematopoietic stem and progenitor cells. In fact, mean pseudotime values among progenitors were highly correlated with mean pseudotime values among myeloid cells but not lymphoid cells (Figure H-J), suggesting that epigenetic reprogramming likely occurs at the level of hematopoietic stem and progenitor cells rather than in mature myeloid cells. In sum, we have proposed a model of epigenetic reprogramming that seeks to explain the acquisition of disease tolerance through repeated exposure as children grow up in a malaria-endemic setting.

## Discussion

In this study, we utilized whole blood transcriptomics and epigenetic profiling at single-cell resolution using EpiTOF to demonstrate that the acquisition of disease tolerance is associated with epigenetic reprogramming of innate immune cells. We observed that symptomatic malaria infections were associate with upregulated interferon signaling and antigen processing/presentation genes and pathways in comparison to an uninfected baseline, similar to prior studies conducted in malaria-naïve adults before and after controlled human malaria infection, and among symptomatic individuals living in malaria-endemic settings compared with uninfected controls^8–10^. In contrast, asymptomatic infections were not associated with any significantly upregulated genes in comparison to an uninfected baseline. These observations are consistent with studies that suggest that prior malaria exposure is associated with a blunted pro-inflammatory transcriptional response in the setting of symptomatic infections^9,11^, but differ somewhat from a smaller study that utilized whole blood gene expression microarray in comparing a set of children with asymptomatic parasitemia with a different set of uninfected controls (n=4/group) that identified genes involving RNA processing and nucleotide binding to differ between asymptomatic parasitemia and uninfected children^10^. Our larger, paired analysis identified that cytokine signaling-related gene sets that were upregulated during symptomatic malaria were downregulated during asymptomatic parasitemia and suggests that regulation of inflammation downstream of TLR signaling contributes to disease tolerance of *P. falciparum* infection.

Our transcriptomic findings motivated an investigation into the epigenetic basis for disease tolerance of malaria. Through multiple EpiTOF experiments performed on separate cohorts, we found that decreased levels of histone H3.3 and increased arginine dimethylation and H3K27 methylation defined an epigenetic signature of innate immune cells that: 1) distinguished asymptomatic parasitemia from symptomatic malaria, 2) correlated with decreased production of cytokines, and 3) predicted the acquisition of disease tolerance in uninfected children. Given these data, we hypothesize that reprogramming of innate immune cell epigenetics plays a causal role in the acquisition of disease tolerance and clinical immunity among Ugandan children repeatedly exposed to malaria.

Our data are further consistent with the hypothesis that repeated exposure to *Plasmodium* epigenetically conditions an innate immune response in favor of disease tolerance^4^. We observed that the relative abundance of immune cells bearing a particular epigenetic signature correlated with future but not past disease tolerance. The importance of temporal directionality in this association suggests that the epigenetic signature associated with disease tolerance likely emerged within the immune cells of Ugandan children over the course of our study as their cumulative parasite exposure increased. Furthermore, pseudotime analyses modelling epigenetic adaptations and the acquisition of disease tolerance revealed both age and recent parasite burden as predictors of epigenetic reprogramming.

The concept of innate immune memory has been defined at the level of disease outcomes through, for example, mouse experiments demonstrating non-specific protection against a myriad of infections following BCG vaccination^14,15,17,18^; however, innate immune memory has also been defined at a cellular level through *in vitro* experiments demonstrating a heightened or refractory response of individual cells to secondary stimulation^21^. Experiments that follow in the form of the latter have clearly demonstrated that an initial stimulus can epigenetically reprogram a cell so as to alter its secondary response^21^. However, this model of innate immune memory may not be relevant in many *in vivo* contexts for the reason that innate immune cells, particularly monocytes, live a rather short life in the circulation. The phenomenon (as we have observed it in this study) that Ugandan children can exhibit disease tolerance and suppression of innate inflammatory responses after months without *Plasmodium* infection (Figure 2B), suggests that innate immune memory is almost certainly not perpetuated by the tolerization of circulating monocytes that persist through multiple infections. Instead, we postulate that epigenetic reprogramming in favor of disease tolerance occurs at the level of stem and progenitor cells in the bone marrow and/or spleen^15^. In support of this hypothesis, the epigenetic signatures that we identified to be associated with disease tolerance and intolerance delineated not only monocytes of symptomatic and asymptomatic children, but also dendritic cells, B cells, and CD8^+^ T cells (Figure 4G). The fact that similar epigenetic signatures were observed across such diverse immune cell populations and were consistent in their associations with disease tolerance suggests mutual descendance from an epigenetically reprogrammed progenitor. Furthermore, we observed a strong correlation between the epigenetic states of hematopoietic progenitors and myeloid cells derived from the same child (Figure 6I). Together, these data support the role of progenitor cells in perpetuating innate immune memory and disease tolerance in the setting of malaria.

There were some limitations of our study. First, while we used multiple cohorts to validate our findings and utilized heterogeneity to improve generalizability, the sample size of these cohorts was relatively small. Further validations should utilize larger cohorts. Second, though we included age and recent parasite burden as independent variables in our cross-sectional models of epigenetic reprogramming, the hypothesis that repeated *Plasmodium* exposure during childhood leads to epigenetic reprogramming should be validated via longitudinal sampling of individual children at multiple timepoints across repeated infections. Third, while our data implicate specific epigenetic marks in disease tolerance, additional studies utilizing methods such as chromatin immunoprecipitation are needed to determine how these markers localize across the genome and affect gene expression. Finally, while our data point to epigenetically reprogrammed progenitors as the mediators of long-term tolerogenic memory, it remains possible that innate immune memory is perpetuated by adaptive immune cells. For example, in a recent study, T cell-induced IFNγ was found to be required to program trained immunity in monocytes following *in vitro* exposure to *Plasmodium falciparum*^29^. In addition, IL-3 is a cytokine that enhances monocyte differentiation, potentiates cytokine storms in sepsis^30^, and contributes to poor disease outcomes associated with blood infections, including malaria^30,31^. Activated T cells and B cells are a primary source of IL-3 in humans^30,31^; and therefore, it remains possible that repeated malaria could condition antigen-specific adaptive immune cells to produce less IL-3 upon activation, leading to less potent activation/priming of innate immune cell responses.

In conclusion, this work demonstrates that the relative abundance of epigenetic markers within immune cells is highly correlative with disease outcomes in *Plasmodium* infection. Moreover, the characteristic epigenetic signatures that we identify do not merely result from either asymptomatic parasitemia or symptomatic malaria; rather, these signatures predict future disease tolerance or intolerance, suggesting a potentially causal role. Furthermore, key markers that defined the epigenetic signature of children primed for tolerance were associated with diminished cytokine responses in peripheral monocytes, demonstrating functional consequences of epigenetic reprogramming. Finally, by connecting age and parasite burden with epigenetics, this study proposes a model to partially explain the acquisition of disease tolerance among Ugandan children through repeated exposed to *Plasmodium*.

## Methods

### Key Resources Table

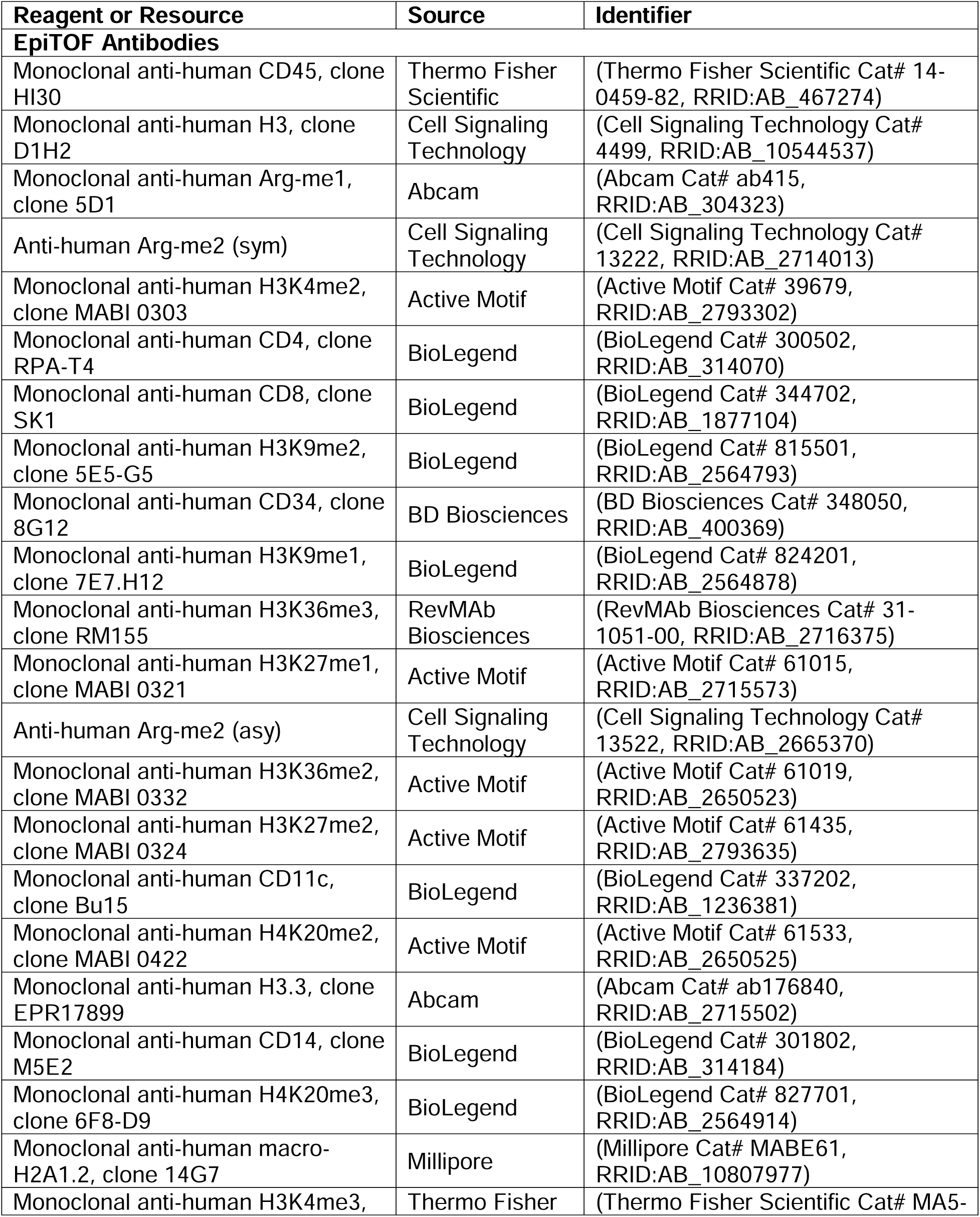
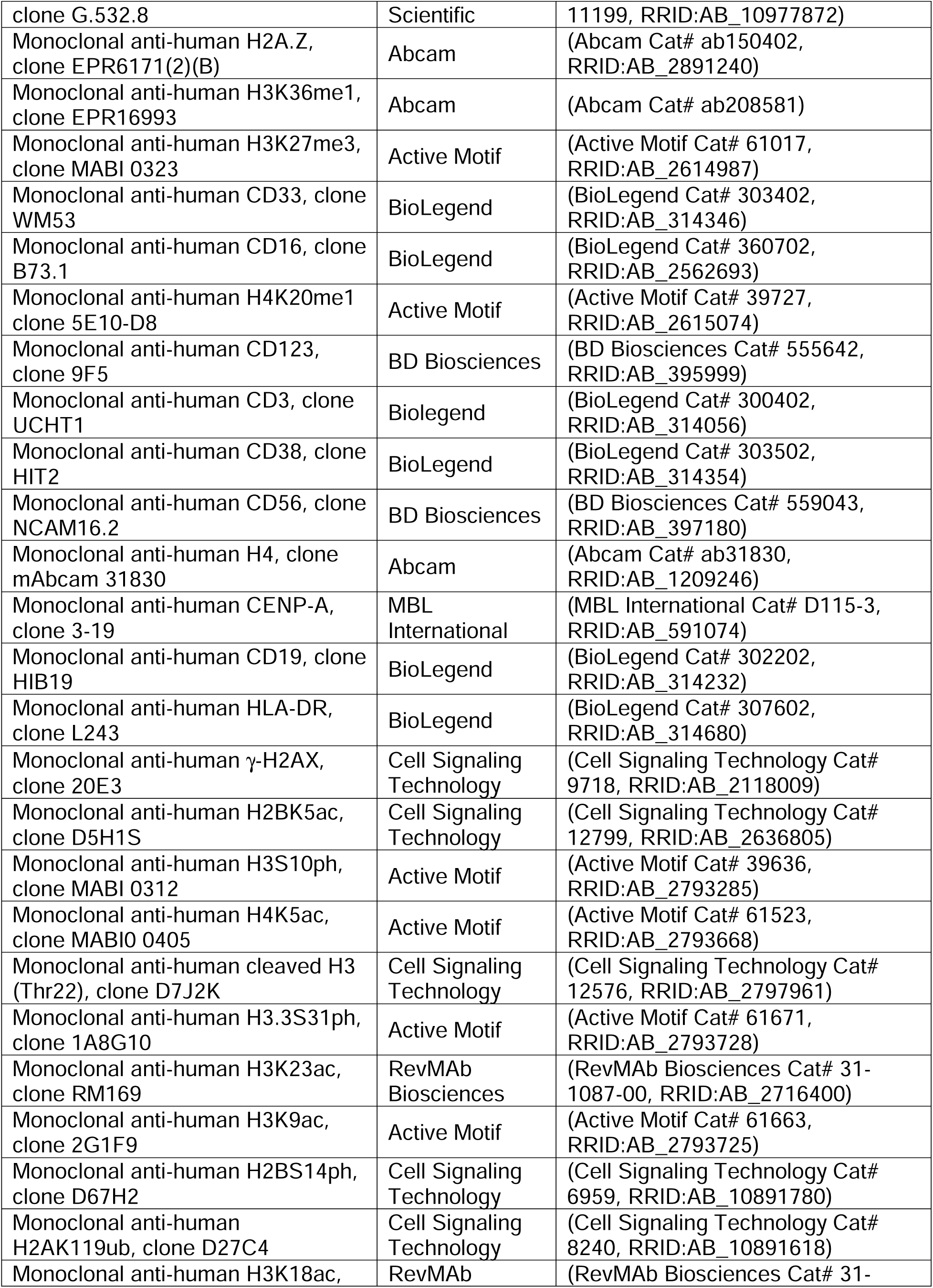
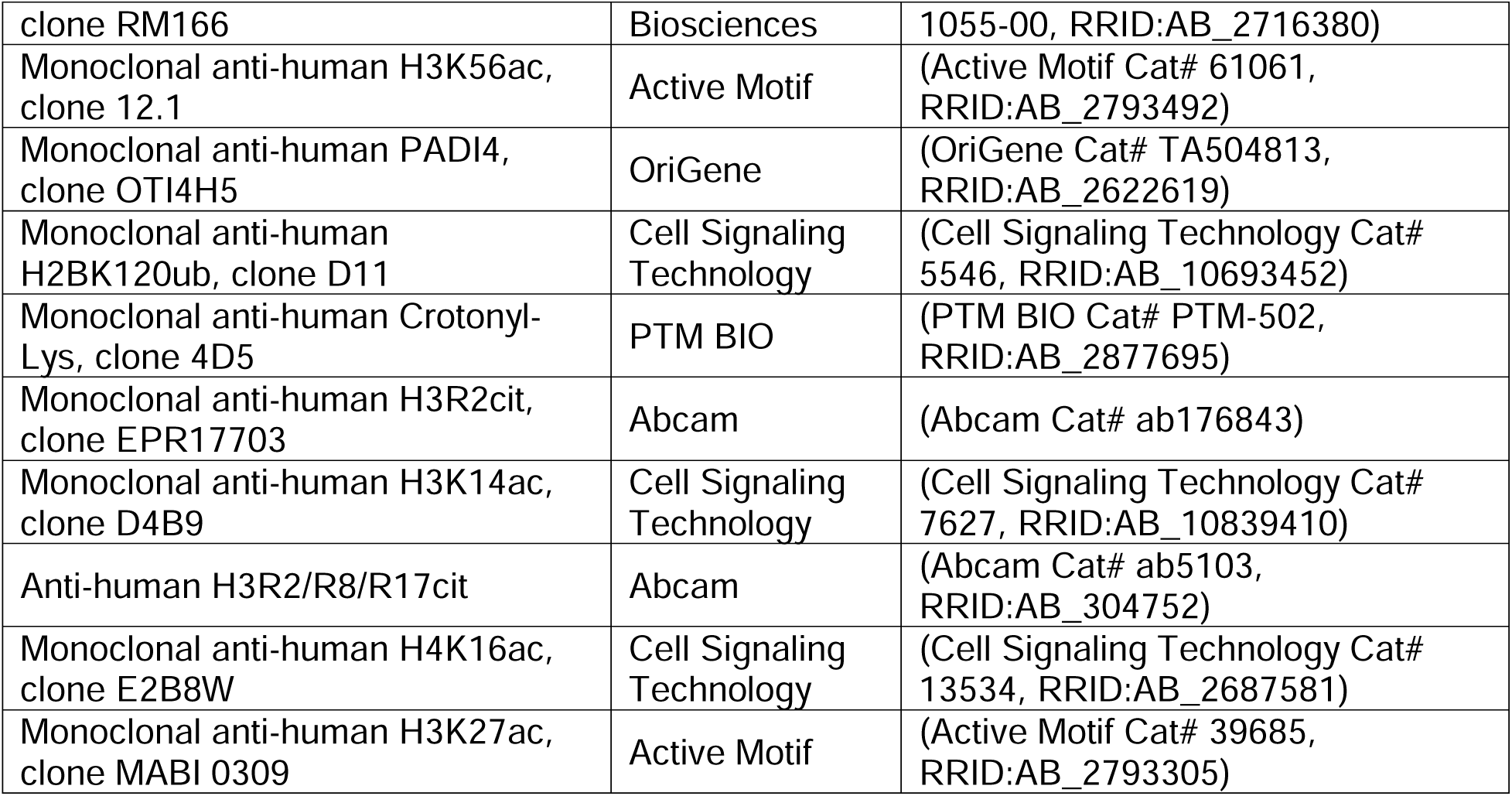

### Study Cohort, Sample Collection, and Processing

Longitudinal samples were obtained from children enrolled in two longitudinal studies in Eastern Uganda. This included children living in Tororo District followed longitudinally through the East African International Centres of Excellence in Malaria Research^32^, and a second cohort of children followed from birth in neighbouring Busia District^33^. In both settings, malaria transmission is high and perennial. In both studies, upon enrollment all participants were given an insecticide treated bed net and followed for all medical care at a dedicated study clinic. Children who presented with a fever (tympanic temperature >38.0 °C) or history of fever in the previous 24 hours had blood obtained by finger prick for a thick smear. If the thick smear was positive for *Plasmodium* parasites, the patient was diagnosed with malaria regardless of parasite density and treated with artemether-lumefantrine. Routine assessments were performed in the study clinic every month, including blood smears and dry blood spots to detect for parasite infection. At select study visits, 3 to 10 milliliters of blood were obtained in EDTA and acid citrate dextrose (ACD) tubes. Blood collected in EDTA tubes (200 μl) was immediately transferred to tubes containing RNA protect. Peripheral blood mononuclear cells (PBMC) were isolated by density gradient centrifugation (Ficoll-Histopaque; GE Life Sciences) from blood collected in ACD tubes, counted, and cryopreserved in liquid nitrogen prior to use.

For RNA Sequencing experiments, paired samples were obtained at two timepoints per subject: when they were uninfected, and when they had either symptomatic or asymptomatic parasitemia. For EpiTOF experiments, paired samples were obtained at three timepoints per subject: when they were uninfected, when they had symptomatic malaria, or when they had asymptomatic parasitemia. The ordering of sampling with respect to disease state differed between individuals.

Written informed consent was obtained from the parent or guardian of all study participants. The study protocols were approved by the Uganda National Council of Science and Technology, the Makerere University School of Medicine Research and Ethics Committee, the University of California, San Francisco Committee on Human Research, and the Institutional Review Boards at Stanford University.

### Whole-blood RNA Sequencing

Whole blood collected in RNA protect was cryopreserved before shipment to Stanford University. For RNA extraction, the mixes of whole blood cells and RNAprotect Cell Reagents were thawed then centrifuged for 5 min at 5000xg. 350 ul QIAGEN Buffer RLT plus was added to dissolve the pellets. The QIAGEN RNeasy Plus mini (QIAGEN, Cat. #74134) kit was used for RNA purification on QIAcube automation (QIAGEN, Cat.# 9002864). RNA samples were eluted in 30ul RNase-free water. All RNAs were checked on a NANODRP1000 and Agilent bioanalyzer 2100 RNA NANO analysis for RNA yield, purity, and integrity. The KAPA mRNA HyperPrep Kits (KK8580) with the IDT for Illumina Dual Index Adapter kit (Cat. #20021454) were used for library preparation per the manufacturer’s protocol. Extra hemoglobin RNA removal and rRNA depletion using the Kapa specific reagents (RiboErase Globin) were added to the protocol. Briefly, mRNA was captured using magnetic oligo-dT beads then RiboErase Globin capture beads were applied. Fragmentation was performed using heat and magnesium. 1st strand cDNA synthesis was completed using random priming. Combined 2nd strand synthesis and A-tailing, adapter ligation, library amplification, and KAPA Pure Beads clean-ups were performed for library preparation. The strand marked with dUTP is not amplified, allowing strand-specific sequencing. All final libraries were checked on Agilent’s bioanalyzer 2100 High Sensitivity DNA Chip. An equal amount of cDNA library from each sample was pooled for sequencing on the Illumina Hiseq 4000 platform. FASTQ files were generated using the bcl2fastq2 Conversion v2.19 tool.

### Epigenetic landscape profiling using cytometry by time of flight (EpiTOF)

Cryopreserved PBMCs were shipped to Stanford University, thawed and incubated in RPMI 1640 media (ThermoFisher) containing 10% FBS (ATCC) at 37°C for 1 hour prior to processing. Cisplatin (ENZO Life Sciences) was added to 10 mM final concentration for viability staining for 5 minutes before quenching with CyTOF Buffer (PBS (ThermoFisher) with 1% BSA (Sigma), 2mM EDTA (Fisher), 0.05% sodium azide). Cells were centrifuged at 400 g for 8 minutes and stained with lanthanide-labeled antibodies against immunophenotypic markers in CyTOF buffer containing Fc receptor blocker (BioLegend) for 30 minutes at room temperature (RT). Following extracellular marker staining, cells were washed 3 times with CyTOF buffer and fixed in 1.6% PFA (Electron Microscopy Sciences) at 1×10^6^ cells/ml for 15 minutes at RT. Cells were centrifuged at 600 g for 5 minutes post-fixation and permeabilized with 1 mL ice-cold methanol (Fisher Scientific) for 20 minutes at 4C. 4 mL of CyTOF buffer was added to stop permeabilization followed by 2 PBS washes. Mass-tag sample barcoding was performed following the manufacturer’s protocol (Fluidigm). Individual samples were then combined and stained with intracellular antibodies in CyTOF buffer containing Fc receptor blocker (BioLegend) overnight at 4C. The following day, cells were washed twice in CyTOF buffer and stained with 250 nM 191/193Ir DNA intercalator (Fluidigm) in PBS with 1.6% PFA for 30 minutes at RT. Cells were washed twice with CyTOF buffer and once with double-deionized water (ddH2O) (ThermoFisher) followed by filtering through 35 mm strainer to remove aggregates. Cells were resuspended in ddH2O containing four element calibration beads (Fluidigm) and analyzed on CyTOF2 (Fluidigm).

### Flow Cytometry

Cryopreserved PBMC samples were thawed in R10 media (RPMI-1640, 10% fetal bovine serum, 1mM L-glutamine, 10mM HEPES, spiked with pen/strep) and allowed to rest for six hours before being stained for surface markers (CD19, CD3, HLA-DR, CD14, CD16, CD11c, CD123). After surface staining, cells were fixed and permeabilized using the kit supplied by ThermoFisher according to the manufacturer’s directions. Intracellular cytokine staining was performed using antibodies specific for IFNγ, TNFα, IL-10, and IL-6. Finally, single cells were analyzed for fluorescence using an Attune NxT flow cytometer. Gating of immune cell populations was performed as shown in Supplementary Figure 2.

## Statistical Methods

### Analysis of Gene Expression Data

The STAR aligner tool was used to trim reads and align them to the reference genome (GRCh38), generating a gene expression count matrix. Differential expression of genes across disease states was analyzed using DeSeq2. Ranked gene lists were used to perform GSEA using Reactome gene sets. Gene sets enriched in children when they had asymptomatic parasitemia versus when they were uninfected (and vice versa) were then assessed for their enrichment in children when they had symptomatic malaria versus when they were uninfected.

### Supervised Analysis of EpiTOF and Flow Cytometry Data

Raw data were pre-processed using FlowJo (FlowJo, LLC) to identify cell events from individual samples by palladium-based mass tags, and to segregate specific immune cell populations by immunophenotypic markers. A detailed gating hierarchy is described in Supplemental Figure 1. Single-cell data for various immune cell subtypes from individual subjects were exported from FlowJo for downstream computational analyses.

The exported Flowjo data were then normalized following previously reported methods^23^. In brief, the value of each histone mark was regressed against the total amount of histones, represented by measured values of H3 and H4. Paired two-tailed T tests were performed to compare the expression of histone markers within specific cell populations across disease states (as in Figure 3A). P values were calculated separately for both cohorts, and then a combined p-value was generated using Fisher’s method. An alpha of 0.01 was used as a threshold of significance to minimize false discovery.

From our flowcytometric data, geometric means of fluorescence intensity (MFI) were calculated for each cytokine and cell population on a per-sample basis. MFI values for stimulated conditions were background subtracted. Linear correlations between histone marker and cytokine expression (z-scored) were performed at the sample-level and permitted random intercepts to account for repeated measurements and variability across disease states.

### Unsupervised Analysis of EpiTOF Data

For unsupervised analyses, weighted sampling of cells was performed to obtain equal numbers of a given cell type across children, across disease states, and across cohorts. UMAP projections and Louvain clustering were performed independently for cohorts 1 and 2 to identify communities of epigenetically related cells. Medians of expression for the 20 markers for each cluster were normalized to those of the other clusters in the same cohort, and then, the Euclidean distance between all clusters was calculated to construct a hierarchy and identify meta clusters.

Fisher’s exact test (two-tailed) was used to compare the frequency of each meta cluster across disease states (as in Figure 4D). In other cases where meta cluster frequencies were calculated on a per-sample basis (as in Figure 4E-G), paired two-tailed t-tests were used to compare these frequencies across disease states. The relative abundance of a meta cluster within a specific immune cell population in children with symptomatic malaria versus asymptomatic parasitemia was calculated using Hedges’ g formula (as in Figure 4G). Regressions involving clinical variables were frequently illustrated as linear relationships with lines of best fit and 95% confidence intervals. For these relationships, Spearman Correlations were also performed with rho values and associated p values displayed.

Clinical variables were calculated as follows: “Malaria Incidence Future” = (# of malaria episodes over the next three years) / 3 years; “Test-Positive Rate Future” = (# of positive blood smears performed over the next three years) / (# of total blood smears performed over the next three years); “log10(GMPD)” = log-base-10 transformation of the geometric mean of all (parasite density values + 1) measured over the next three years; “NMF Incidence Future” = (# of non-malarial fevers observed over the next three years) / 3 years; “Future Risk of Malaria if Parasitemic” = Malaria Incidence Future / (Test-Positive Rate Future x 365); “Past Risk of Malaria if Parasitemic” = Malaria Incidence Past / (Test-Positive Rate Past x 365).

### Pseudotime Analysis of EpiTOF Data

Tspace was used to perform trajectory inference with the same data used in our other unsupervised analyses. Standard algorithm parameters were used as recommended by the developers. The expression of epigenetic markers over pseudotime was fit using an 8th-degree polynomial function (Figure 6C). Correlations with clinical variables were performed as described above except for the multiple regression of mean pseudotime on “Age” and “GMPD Last 90 Days,” which utilized standardized variables.

## Data Availability

Whole-blood RNAseq data for this study have been deposited on the National Center for Biotechnology Information’s Gene Expression Omnibus under the accession number: ––. EpiTOF data have been made available on ImmPort.

## Acknowledgements

We would like to acknowledge and thank the Human Immune Monitoring Center (HIMC) at Stanford Medicine for helping perform the RNAseq experiments in this study. We would also like to acknowledge Kattria van der Ploeg for providing her valuable feedback during the completion of this manuscript.

## Author Contributions

J.N., B.P., P.K., and P.J. conceptualized the manuscript. X.J. supervised RNAseq experiments. P.U. supervised EpiTOF experiments. R.J., R.K., F.N., K.M., and M.K. oversaw the clinical studies and/or collected/processed patient samples. M.K., M.F., G.D. provided samples for the study. B.P., P.K., and P.J. designed the experiments and advised with analyses. J.N. and M.T. performed the data analyses. J.N. and P.J. wrote the manuscript. All authors have read and support the findings presented in the manuscript.

## Declaration of Interests

The authors declare no competing interests.

## Notes

### Competing Interest Statement

The authors have declared no competing interest.

