## Supplementary Figure 1 for "Disease Tolerance Acquired Through Repeated *Plasmodium* Infection Involves Epigenetic Reprogramming of Innate Immune Cells"

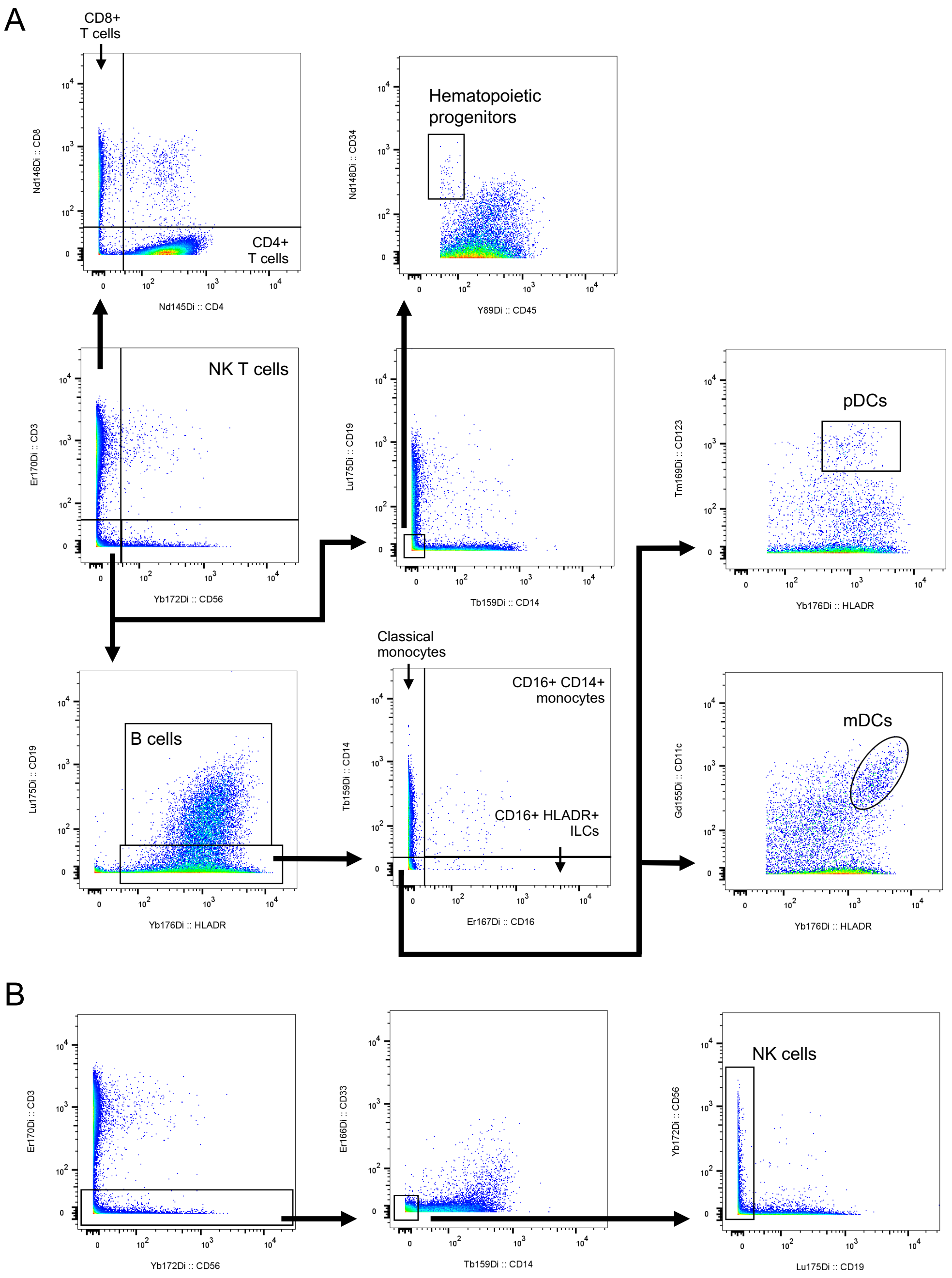


***Supplementary Figure 1**. Gating strategy for identifying surface marker defined cell populations from EpiTOF experiments.* (A) Gating of annotated cell populations distinct from NK cells. (B) Gating of NK cells.
