## Supplementary Figure 2 for "Disease Tolerance Acquired Through Repeated *Plasmodium* Infection Involves Epigenetic Reprogramming of Innate Immune Cells"

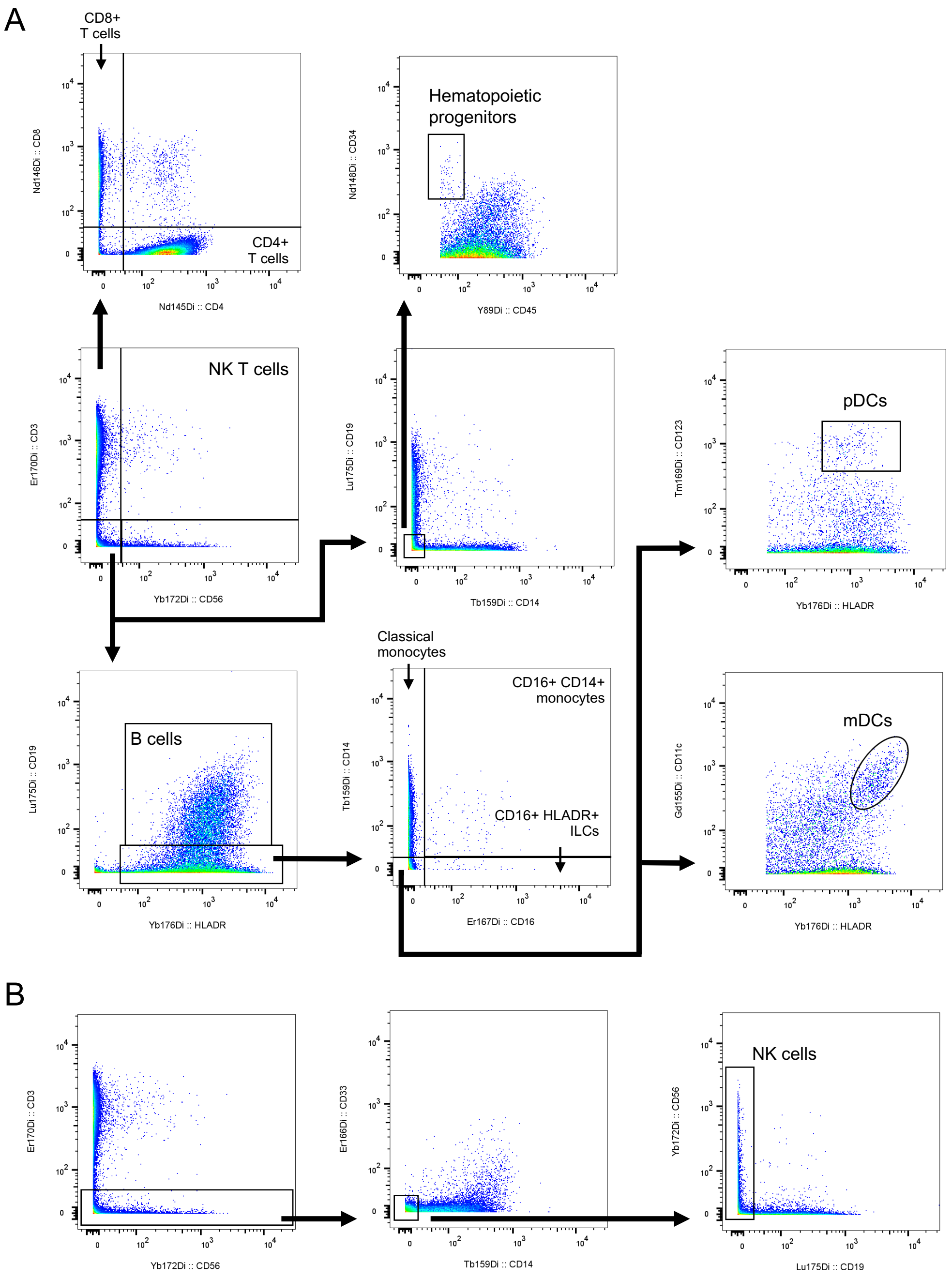


***Supplementary Figure 1**. Gating strategy for identifying surface marker defined cell populations from EpiTOF experiments.* (A) Gating of annotated cell populations distinct from NK cells. (B) Gating of NK cells.

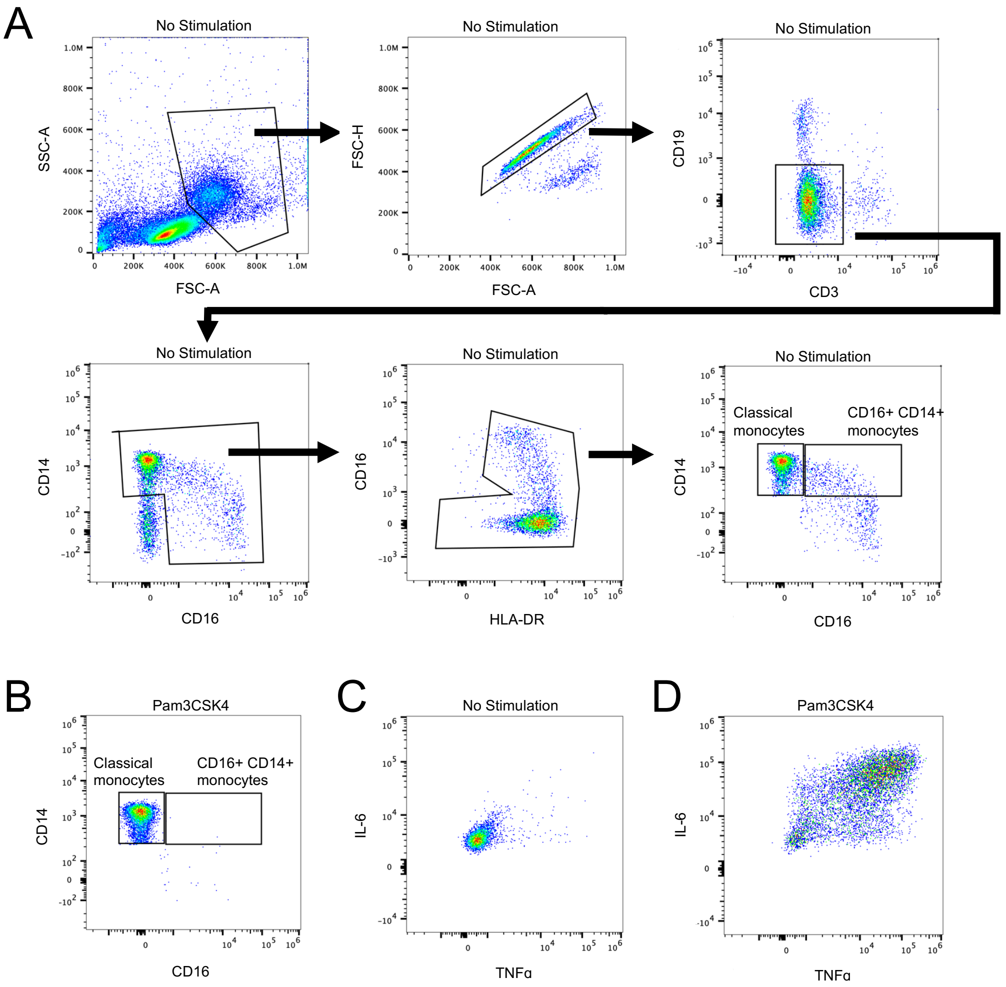


***Supplementary Figure 2****. Gating strategy for monocytes stimulated and analyzed by flow cytometry.* (A) Gating of classical and CD16+ monocytes. (B) Stimulation with Pam3CSK4 kills CD16+ monocytes but not classical monocytes. (C-D) Representative flow plots showing TNFα and IL-6 expression of classical monocytes after no stimulation (C) or Pam3CSK4 stimulation (D).
